# Cross-protocol comparison of iPSC-microglia reveals hypofunction contributes to neuronal vulnerability and synaptic alterations in the *MAPT*-S305N model of frontotemporal dementia

**DOI:** 10.64898/2026.06.30.735652

**Authors:** D Vasoya, LK Keavey, C Levit, T Watzeels, S Heron, J Cholewa-Waclaw, O Dando, R Mancuso, KR Bowles

**Author notes:** Corresponding author: Contact, Kathryn R Bowles.

## Abstract

Progressive and chronic neuroinflammation is associated with numerous neurodegenerative diseases, including primary tauopathies such as frontotemporal dementia and progressive supranuclear palsy. Unlike Alzheimer’s disease, there is no clear genetic association implicating microglial dysfunction as a primary driver of tauopathy. As such, the contributions of microglia to tauopathy pathogenesis have been less well defined. Here, we explore the cell autonomous effects of the pathogenic *MAPT*-S305N variant on microglial function, across two distinct iPSC-microglia protocols, followed by examination of the non-cell autonomous effects of microglial *MAPT* genotype on neuronal health and function. We find that different protocols produce cells of equivalent microglial identity, but result in microglia in different functional states, thereby influencing reactivity and detectable phenotypes. Regardless, across both protocols we find that *MAPT*-S305N induces microglial hypoactivity, evidenced by impaired phagocytosis, reduced cytokine release and diminished regulation of synaptic function. We conclude that microglial hypoactivity may be an early event in disease pathogenesis, where *MAPT* mutation microglia fail to adequately respond to pathogenic stimuli, thereby contributing to subsequent neuronal vulnerability and susceptibility. Further studies are required to understand how and when this initial hypoactive state may switch to a toxic pro-inflammatory state, and whether early detection and correction may be of therapeutic value.

## INTRODUCTION

Neuroinflammation and microglial dysfunction are understood to be primary drivers of several neurodegenerative diseases, including Alzheimer’s disease (AD)^1^ and Parkinson’s disease (PD)^2^, supported by observations that common variants in microglia-specific genes confer increased risk^3,4^. However, the contribution of neuroinflammation to primary tauopathies, such as progressive supranuclear palsy (PSP) and frontotemporal dementia (FTD-tau) is less clear; to date, there are no genome-wide association studies (GWAS) that implicate microglial function in PSP or FTD-tau risk, with microglial enhancers and promoters estimated to be the least impacted by risk-associated variants compared to neurons, astrocytes or oligodendroglia^5^. Regardless, multiple streams of evidence still point to a role for microglia function and neuroinflammation in modulating disease-associated phenotypes. For example, assessment of a PET tracer for neuroinflammation indicates that familial FTD and PSP progression correlates with tracer positivity to a similar extent as fluortaucipir, a tracer sensitive to tau pathology^6^, suggestive of a role for neuroinflammation in the progression and severity of disease.

Targeted studies both *in vivo* and *in vitro* are also indicative of a neuroinflammatory effect of pathogenic tau^7–10^, while functional impairments have also been observed in tauopathy model microglia^11,12^. Interestingly, microglia derived from *MAPT*-P301S mice were observed to aberrantly phagocytose live neurons, and were subsequently less able to internalise exogenously applied tau seeds^12^, possibly contributing to neurodegeneration and tau spread. Similarly, microglial *APOE* genotype has been shown to modulate the extent of tau pathology in a similar *MAPT*-P301S mouse model^13^, indicating that microglial function does indeed have the capacity to modulate tauopathy disease progression.

However, questions remain regarding whether microglia may be passive responders to neuronal dysfunction and pathological tau accumulation, or whether there may be a cell autonomous effect of pathological *MAPT* expression that modulates microglial function and actively contributes to disease progression. The latter has remained largely unexplored, as *MAPT* expression in microglia is minimal, and tau pathology in this cell type is rare. However, a recent report demonstrated not only detectable *MAPT* expression in iPSC-derived microglia, but also cell-autonomous effects of the *MAPT*-IVS10+16 mutation on microglial function^11^.

Given these observations, we decided to explore both the cell-autonomous and non-cell autonomous impact of the *MAPT*-S305N splicing mutation on microglial function and neuronal health in iPSC models, in order to understand the nature of microglial contributions to cellular dysfunction in primary tauopathies. However, before we could embark on this effort, we needed to select an appropriate microglia differentiation protocol to use for the project. Numerous microglia differentiation protocols have been published, collated and reviewed elsewhere^14^, yet two of the most popular and widely adopted protocols: McQuade *et al*^15^ and Haenseler *et al*^16^, both published in 2017, have never been directly compared to each other. Curiously, adoption of each protocol appears to follow a geographical divide across the Atlantic Ocean, with McQuade *et al* (hereafter referred to as the “HPC” protocol) being widely used in the USA, while Haenseler *et al* (“Factory” protocol) is more popular in the UK and Europe, most likely reflecting the region in which each protocol was originally developed. Importantly, both approaches have been thoroughly characterised as microglia-like cells, and have been successfully independently used by many groups. Even so, we could not determine from published data which approach would be best suited for our scientific question. Therefore, rather than selecting one protocol, we decided to conduct our experiments in microglia derived from both protocols in parallel. This has allowed us to not only compare and characterise microglia across protocols in cells derived from the same iPSC lines, but also to determine which aspects of *MAPT*-mutation-induced microglia dysfunction are most robust and reproducible across models.

Here, we report that each protocol results in microglia with equivalent levels of appropriate identity markers, however they seemingly produce microglia in different states of function or activation. Subsequently, the starting microglial state defined the phenotypes that were detected in *MAPT*-S305N microglia compared to their isogenic controls. Nevertheless, we conclude that the *MAPT*-S305N mutation has a cell autonomous functional impact on microglia by suppressing reactivity to stimuli. Subsequently, this impairment is detrimental to synaptic regulation and function due to an inability to appropriately respond to pathogenic environments. As such, we propose microglia may contribute to early disease pathogenesis by failing to adequately support neuronal health through inaction, rather than by overactivation and inflammation.

## METHODS

### iPSC lines

Two fc and matched wild type (WT) control iPSC lines were used throughout the study; 300.12-NWBB7 (WT/WT) and 300.12-NW1C7 (S305N/WT), and F13505-S2A3 (WT/WT) and F13505-NDG11 (S305N/S305N). The generation of these lines has been previously described^17^, and lines are available from the National Centralized Repository for Alzheimer’s Disease (NCRAD; https://ncrad.iu.edu/) and the Tau Consortium via the Neural Stem Cell Institute (NSCI; http://neuralsci.org/tau).

iPSCs were maintained in StemFlex media (ThermoFisher Sci) supplemented with 1% antibiotic-antimycotic (ThermoFisher Sci) on Matrigel (Corning)-coated plates. Lines were passaged weekly using ReLeSR (StemCell Technologies). All cell cultures were maintained at 37C:C with 5% CO_2_ in a humid environment.

### Microglia differentiation

#### Factories

“Factory” protocol microglia were generated as described in Haenseler et al^16^. iPSCs were dissociated using StemPro Accutase (ThermoFisher Scientific), followed by a viability check and cell count using Trypan Blue visualised on a Countess (ThermoFisher Sci). Three wells of a 6-well plate of 80% confluent iPSCs were collected and pelleted by centrifugation at 70xg for 3 minutes, before resuspension in 10mL EB buffer (StemFlex, 50ng/mL BMP4 (Qkine), 50ng/mL VEGF (Qkine), 20ng/mL SCF (ThermoFisher Sci)) supplemented with 10µM Y27632 (Fisher Scientific). The cell suspension was divided equally across all wells of Nunclon Sphera U-bottom 96-well plate (ThermoFisher Sci), which was then centrifuged for 3 minutes at 100xg to collect cells into an embryoid body. After 48 hours, cells underwent a 50% media change with fresh EB media, without Y27632 supplementation. Following another 48 hours, EBs were collected and transferred into Factory media (X-VIVO15 (Lonza), 1% Glutamax (ThermoFisher Sci), 1% antibiotic-antimycotic, 0.1% β-Mercaptoethanol (Fisher Scientific), 100ng/mL M-CSF (Qkine), 25ng/mL IL-3 (Qkine)) in Matrigel-coated 6-well plates. Two thirds of the media was replaced every 5 days with fresh Factory media for three weeks. At this point, monocyte progenitor cells were visible in the supernatant. Progenitors were harvested by collecting the supernatant and centrifuging for 5 minutes at 400xg. The first harvest was always discarded, and the second harvest was used for experiments. Collected progenitors were resuspended in Differentiation media (Advanced DMEM/F12 (ThermoFisher Sci), 1% Glutamax, 1% antibiotic-antimycotic, 100ng/mL IL-34 (ThermoFisher Sci), 10ng/mL GM-CSF (ThermoFisher Sci), 25ng/mL M-CSF (Qkine), 50ng/mL TGFβ1 (ThermoFisher Sci)) and plated into the required final format; 0.75 x 10^6^ per well for 6-well plates (RNA-seq, cytokine release), 1.5 x 10^4^ per well for 96-well plates (phagocytosis assays) and 3 x 10^4^ per chamber for 8-chamber imaging slides (immunofluorescence). Cells were matured for 10 days in Differentiation media before use, with 50% media changes every 3-4 days. While Factories can continue to be maintained for a long period with multiple rounds of harvesting, for this study we collected only the second harvest for all assays. Each differentiation batch therefore refers to independent differentiations initiated from iPSC, not to different harvests derived from one factory culture.

#### HPCs

“HPC” protocol microglia were generated as described in McQuade et al^15^. iPSCs were dissociated using ReLeSR and transferred at low density into Matrigel-coated 6-well plates. Multiple split densities (1:20, 1:40, 1:60) were used for each line each time, as the efficiency of differentiation was sensitive to the starting number of cells and variable between lines and batches. After 24 hours, media was replaced with HPC-A media from the StemDiff Hematopoietic Kit (StemCell Technologies). A 50% media change with fresh HPC-A was performed after 48 hours, followed by a full media change to HPC-B the next day. 1mL fresh HPC-B was added to cultures every 2 days for one week. The first collection of hematopoietic cells (HPCs) was then performed by collecting the supernatant. The first collection was discarded as it typically contained a high proportion of debris and dead cells. The second collection was performed after 2 more days in culture, and cells were pelleted by centrifugation at 300xg for 5 minutes. The collected HPCs were transferred into Differentiation media (DMEM/F12 (ThermoFisher Sci), 2x B27 (ThermoFisher Sci), 0.5x N2 (ThermoFisher Sci), 1% Glutamax, 1% non-essential amino acids (ThermoFisher Sci), 400µM monothioglycerol (Sigma Aldrich), 5µg/mL human insulin (Sigma Aldrich), 1% antibiotic-antimycotic, 100ng/mL IL-34, 25ng/mL M-CSF, 50ng/mL TGFβ1)) and seeded into Matrigel-coated 6-well plates at a density of 1 x 10^5^ cells per well. 1mL of fresh Differentiation media was added every 2 days, until the volume in the well was too high. At this point, media and cells in suspension were collected and spun at 300xg for 5 minutes to collect the cells. Cells were resuspended in 1mL fresh Differentiation media and returned to culture with 1mL conditioned media. Culture in differentiation media continued for 24 days, after which the media was supplemented with maturation factors; 100ng/mL CD200 (StemCell Technologies) and 100ng/mL CX3CL1 (ThermoFisher Sci). After 2 more days, an additional 1mL of Differentiation media plus maturation factors were added to cells. The next day, microglia were collected and plated in the required format (at the same densities as described for Factories), and used for assessment after an additional 24 hours.

#### Phagocytosis assays

Microglia were treated with 10ng/mL pHrodo-conjugated E.Coli particles and were immediately imaged using the IncuCyte live cell imaging system (EssenBio). As a negative control, microglia were pre-treated with 10µM cytochalasin D (Cambridge Bio) for 30 minutes to inhibit phagocytosis, prior to E.Coli treatment. Three fields of view per well were imaged every hour for 24 hours, using a 20x objective. Image analysis was carried out in the IncuCyte software for automated measurement of pHrodo positive area, normalised to cell area (determined by brightfield imaging).

#### Synaptosome preparation

Cortical organoids were generated from F13505-S2A3 and F13505-NDG11 iPSC lines using an established and previously described protocol^18^. 30 organoids per line were collected at 6 months of differentiation, and synaptosomes were isolated using the Syn-PER synaptic protein extraction reagent (ThermoFisher Sci). Organoids were homogenized in Syn-Per reagent in a Dounce homogenizer for 10 strokes, then centrifuged at 1,200xg for 10 minutes at 4C:C. The pellet was discarded, and the supernatant centrifuged again for 20 minutes at 15,000xg at 4C:C. The supernatant was retained as the cytosolic fraction, and the pellet was resuspended in Syn-Per reagent as the synaptic fraction.

#### Western blot

Synaptosome identity was confirmed by western blot; briefly, cytosolic and synaptic fractions were incubated with 1 x reducing agent and 1 x LDS sample buffer (both ThermoFisher Sci) at 70C:C for 10 minutes, before immediately being loaded onto a Bolt 4-12% Bis-Tris Plus gel in 1x MES buffer (ThermoFisher Sci). Electrophoresis was carried out at 220V for 22 minutes, before blotting onto nitrocellulose membranes using the iBlot system (ThermoFisher Sci). The membrane was blocked for a minimum of 30 minutes at room temperature in 5% milk in PBS-T. Primary antibodies were prepared in 5% milk as follows; Homer1 (1:1000, Synaptic Systems), Da9 (total tau, 1:500, a kind gift from Drs. Jeremy Koppel and Peter Davies) and GAPDH (1:10,000, Cell Signaling Technology). Antibodies were incubated with membranes overnight at 4C:C on a rocker, then washed 3 x with PBS-T. Membranes were then incubated with secondary antibodies prepared in 5% milk at room temperature for an hour; IRdye 680RD anti-Rabbit (1:10,000) and IRDye 800CW anti-mouse (1:10,000; both LICORbio). Following three additional washes, membranes were imaged on the LICOR Odyssey CLx imager. For re-staining, membranes were stripped with Restore Fluorescent Western Blot Stripping buffer (ThermoFisher Sci) for 30 minutes at room temperature, before a 5 minute wash in PBS and re-blocking in 5% milk.

#### pHrodo labelling

Synaptosome preparations were pelleted by centrifugation at 15,000xg for 3 minutes at 4C:C, and resuspended in 0.1M sodium bicarbonate buffer in 0.1% BSA (pH 8.4). pHrodo ester (ThermoFisher Sci) resuspended in 150µl DMSO (Sigma Aldrich) was added to synaptosomes at a 1:10 dilution, mixed, and incubated in the dark for 60 minutes. PBS was added to the labelled sample, and centrifuged for 3 minutes at 15,000xg, after which the supernatant was discarded. The PBS wash was repeated two more times, and the resulting labelled synaptosomes were resuspended in PBS to give a concentration of 10µg/µl protein. Synaptosome preparations were added to microglia at 10µg/well, and imaged and analysed in the same way as for E.Coli described above.

### Stimulation

To assess the effect of protocol and mutation on microglial responses to stimuli, cells were exposed to either 20ng/mL IL-4 (Qkine) or 10ng/mL lipopolysaccharide (LPS; ThermoFisher Sci) for 24 hours. The equivalent volume of sterile water was used as the carrier control.

### Inflammation marker profiling

Cytokine and chemokine secretion was initially screened using Proteome Profiler Human Cytokine Array kit (R&D systems). Conditioned microglia media was collected after 24 hours incubation with either IL-4, LPS or carrier control. Media was centrifuged at 1,000xg for 5 minutes to pellet remaining cells and debris. Cytokine arrays were carried out according to the manufacturer’s protocol, using 1mL of conditioned media from each sample. Densitometry was carried out in ImageJ 1.53q (http://imagej.nih.gov/ij).

Select cytokines were validated by ELISAs, following manufacturer’s instructions; MIF, IL-6 and IL-21 (all ThermoFisher Sci).

### Immunofluorescence

For microglia monocultures, immunofluorescence was carried out in 8-well ibiTreat µ-Slides (Ibidi), and co-cultures used 96-well square µ-Plates (Ibidi). Cells were permeabilized and blocked in 0.1% Triton X-100 and 1% BSA in PBS for 30 minutes at room temperature with gentle rocking, followed by 3 x washes with PBS. Primary antibodies were prepared in 1% BSA in PBS, and incubated on cells overnight at 4C:C, followed by 3 more PBS washes and incubation with the appropriate secondary antibodies for 2 hours at room temperature. Secondary antibodies were then removed, and cells were incubated for 10 minutes at room temperature with 300nM DAPI, prepared in PBS. A final three PBS washes were conducted, and cells were stored in PBS at 4C:C until imaging.

For monoculture microglia, the following primary antibodies were used: IBA1 (MA5-38266, 1:100 - ThermoFisher Sci), P2Y12 (4H5L19, 1:500 – ThermoFisher Sci), iNOS (MA5-17139, 1:200 – ThermoFisher Sci), ARG1 (PA5-29645, 1:200 – ThermoFisher Sci) and CD68 (ab289671, 1:500 – Abcam).

For co-cultures, the following additional primary antibodies were used: TMEM176B (PA5-112751, 1:100 – ThermoFisher Sci), Synapsin I (106 009, 1:500 - Synaptic Systems), GFAP (3670S, 1:500 – Cell Signaling Technology) and B-III-tubulin (PA5-85639, 1:100 – ThermoFisher Sci).

Secondary antibodies listed below were all AlexaFluors and were used at 1:100, and were purchased from ThermoFisher Sci: goat-anti-Rat 488, goat-anti-mouse 568, goat-anti-mouse 647, goat-anti-Rabbit 568, goat-anti-chicken 647, goat-anti-guinea pig 568, donkey-anti-goat 647.

Slides and plates were imaged on the Opera Phenix Plus high-content spinning disc confocal microscope (Revvity) using a 40x water objective (NA 1.1). A Z-stack of 5 slices spaced at 2.5µm were taken, with 20 fields of view automatically obtained from each well. The microscopist was blinded to genotype or culture condition.

### Image analysis

Microglia monoculture images were analysed using Harmony software v.5.2 (Revvity). Nuclei were segmented using DAPI, and the morphological characteristics based on IBA1 signal were measured. Fluorescence intensities for each channel were calculated and averaged over each field of view within each replicate well.

For co-cultures, images were processed in ImageJ (Fiji) v.2.3.0. For co-localisation analysis, each channel was automatically thresholded using the Otsu algorithm and converted to a mask. Channel overlaps between Synapsin I and IBA1 were determined by the imageCalculator function, and the area of overlap was normalised to IBA1 area for each field of view. For intensity analysis, following thresholding as described above, mean TMEM176B fluorescence intensity within IBA1 positive areas was measured. Data were averaged across all fields of view within each replicate well.

### RNA sequencing

For transcriptomic assessment, microglia were collected and pelleted, followed by RNA extraction using the RNeasy kit (Qiagen). RNA was submitted to Novogene for mRNA library preparation with polyA enrichment, followed by 150bp paired end sequencing on the Novaseq X Plus.

Samples were sequenced to a depth of approximately 37 million reads. The reads were mapped to the primary assembly of the human (GRCh38) reference genome contained in Ensembl release 113, using the STAR RNA-seq aligner, v.2.7.9a^19^. Tables of per-gene read counts were generated from the mapped reads with featureCounts, v.2.0.2^20^. Differential gene expression was performed in R using DESeq2, v.1.36.0^21^. Gene ontology enrichment analysis was performed using topGO version 2.48.0^22^, and gene set testing was performed using Camera^23^ from the R package limma^24^, v3.52.4, using gene sets from the Molecular Signatures Database, v.7.5.1 (https://www.gsea-msigdb.org/gsea/msigdb/).

### Comparison to xenografted human microglia states

Transcriptional reference profiles for in vivo human microglia states were taken from the single-cell xenotransplantation dataset of Mancuso et al^25^. Cells were grouped by their annotated microglial state, and a per-state pseudobulk profile was generated by summing raw counts across all cells of each state (Seurat v5.5.0 AggregateExpression), followed by log-normalization (Seurat NormalizeData). For marker-gene analyses, state-defining markers were obtained with Seurat FindAllMarkers on the log-normalized single-cell data (min.pct = 0.1), retaining positive markers per state at Bonferroni adjusted P < 0.05 and avg log2FC > 0.

Factory, HPC and the per-state xenograft pseudobulk profiles were reduced to the set of genes shared across all three datasets (17,984 genes). Pairwise sample-to-sample similarity was computed across all averaged profiles using Pearson correlation. Correlation heatmaps were generated with pheatmap v1.0.13.

Differential expression between Factory and HPC protocol microglia was performed on the wild-type, untreated bulk samples. FPKM values were log2-transformed (log2(FPKM + 1)) and lowly expressed genes were removed (mean FPKM > 1). A linear model was fitted with protocol as the sole factor (Factory protocol as the reference level) using limma v3.68.4^24^ with the trend option (lmFit, eBayes(trend = TRUE)). Genes significantly up in HPC protocol (positive log2FC, Benjamini-Hochberg adjusted P < 0.05) and significantly up in Factory protocol (negative log2FC, Benjamini-Hochberg adjusted P < 0.05) microglia defined the two protocol-characteristic gene sets.

For each xenograft microglia state, the set of positive state markers (adjusted P < 0.05, avg log2FC > 0; from FindAllMarkers above) was intersected with the protocol-characteristic gene sets, restricted to the genes shared between the bulk and single-cell data, and used to quantify, for each protocol, how many of each state’s markers were recovered and what fraction of that state’s marker set this represented.

### Neuron-microglia co-culture

Neurons were differentiated from neural progenitor cells (NPCs) derived from the same isogenic pairs of iPSC lines, using previously established protocols^26,27^. Briefly, NPCs were derived from iPSC using Neural Induction Media (NIM; StemCell Technologies), supplemented with 10nM SB431542 and 1nM LDN-193189 (both StemCell Technologies). Rosettes were selected using Rosette Selection Reagent (StemCell Technologies), and cultured for an additional week in NIM. Cells were then transitioned to NPC media (DMEM/F12, 1x B27, 1x N2, 1% antibiotic-antimycotic, 20ng/mL FGF2 (Qkine) for expansion, followed by isolation of CD271-negative and CD133-positive cells by magnetic activated cell sorting (MACS; all reagents from Miltenyi) to produce a homogenous population of NPCs. NPCs were maintained in NPC media and passaged using Accutase, then seeded into the required format at a density of 2.5 x 10^4^ per well in a 96 well plate for immunofluorescence or multi-electrode array (MEA), or 2 x 10^5^ per well in a 6-well plate for single cell sequencing. Cells were then maintained in Neuron media (BrainPhys (StemCell Technologies), 1x B27, 20ng/mL BDNF (Qkine), 20ng/mL GDNF (Proteintech), 250µg/mL cyclic-AMP (Sigma Aldrich), 200µM ascorbic acid (Sigma Aldrich), 1% antibiotic-antimycotic) for 4 weeks, with a full media change every two – three days.

Cultures were timed so that Factory progenitors could be harvested when neuronal cultures reached 4 weeks of differentiation from NPCs. Progenitors were collected by centrifugation as described above, and resuspended in Co-Culture media (Advanced DMEM/F12, 1x N2, 1x Glutamax, 1% antibiotic-antimycotic, 100ng/mL IL-34, 10ng/mL GM-CSF). Neuronal media was removed from neuron cultures, and replaced with progenitors in Co-Culture media, at the same density as NPCs were originally seeded. Cells were maintained together in co-culture for an additional 2 weeks prior to collection and processing, with 50% media changes every other day.

### Single cell sequencing

Co-cultures were gently dissociated with a Papain Dissociation Kit (Worthington Biochemical) for preparation for single cell sequencing, as described previously^28^. Briefly, media was aspirated from cell cultures, and replaced with 1mL prepared papain supplemented with DNase I. Cells were incubated at 37C:C for 90 minutes, with trituration every 15-30 minutes to aid with dissociation. Single cell suspensions were collected through a 40µm filter, pelleted and fixed using the Chromium Next GEM Single Cell Fixed RNA Sample Preparation Kit (10x Genomics). Cells underwent library preparation using GEM-X Flex v2 chemistry at the University of Edinburgh Single Cell Sequencing facility. Libraries were sequenced at the University of Edinburgh Clinical Research Facility, on the NextSeq 2000 platform (Illumina). Basecall data was demultiplexed using the CellRanger *mkfastq* wrapper.

Resulting fastq files underwent quality control and alignment to the human hg38 genome via the 10x Genomics Cloud using the CellRanger analysis pipeline with default settings. The resulting matrix files were imported and processed in Seurat v5.1.0^29–32^. Data were filtered for cells with > 200 and < 10,000 reads, and with mitochondrial gene expression rates below 5%. Data were normalised using SCTransform^33^ and integrated using RPCA, accounting for mitochondrial gene percentage, number of genes and number of reads per cell. Principal components analysis was carried out using the top 3000 most variable genes, and data reduction was performed with UMAP^34^. Broad cell types were determined by positivity for *MAPT* (neurons), *GFAP* (astrocytes) and *MS4A4A* (microglia). Neurons and microglia were individually subset from the full dataset for differential gene expression analysis, following re-scaling and reduction. Differential gene expression was carried out on the raw count data between conditions using the default *FindMarkers* command within Seurat (Wilcoxon rank sum test). The same parameters were kept for all comparisons. Significantly differentially expressed genes (adjusted p.value < 0.05) were submitted to Gene Ontology (GO) pathway analysis in g:profiler^35^.

### Multi-electrode array

For measurement of neuronal network activity, neuron-microglia co-cultures were cultured on 48-well Lumos MEA plates (Axion Biosystems) at 5 x 10^4^ neurons and 5 x 10^4^ microglia seeded per well. Activity was recorded for 5 minutes on the Axion Maestro Pro (Axion Biosystems) using default neural real time spontaneous activity settings, following an initial 1-minute equilibration period. During recording, cultures were maintained at 37C:C with 5% CO_2._ Activity data was processed using the Axion AxiS software with default Neural Activity settings.

### Statistical approach

RNA sequencing and single cell RNA sequencing data were analysed as described above. Quantification of immunofluorescence data was carried out per field of view, and averaged over each replicate well. Data was analysed using a One-way or Mixed-Effects ANOVA and Tukey’s multiple comparisons test to account for different genotypes, conditions and microglia protocols being assessed, while accounting for relatedness of samples. Quantification of phagocytosis data was analysed using a Multiple Linear Regression with Holm-Šídák’s multiple comparisons test to compare the slopes under each condition. Absorbance values obtained from ELISAs were aligned to standard curves using a Four Parameter Logistic (4PL) Regression, and comparisons between groups were conducted using a Mixed-Effects ANOVA with Tukey’s multiple comparisons test. All statistical analyses were carried out either in GraphPad Prism 10, or R v4.4.0. All experiments were conducted on two isogenic pairs of *MAPT*-S305N and corrected control iPSC lines, using three independent rounds of differentiation, unless otherwise stated in figure legends. Specific statistical tests and N for individual assays are described in the relevant figure legends.

## RESULTS

### Factory and HPC iPSC protocols produce microglia with distinct morphologies and transcriptomic profiles

All iPSC differentiation protocols strive to accurately recapitulate cellular identities and states observed *in vivo* as closely as possible. While both Factory and HPC protocols have independently been benchmarked against primary human microglia, and have been shown to be positive for relevant microglial identity makers^15,16^, they have not yet been directly compared to each other. As such, we generated iPSC-microglia from two isogenic pairs of *MAPT*-S305N mutation iPSC lines^17^ using both protocols in parallel, in order to determine whether mutation-associated phenotypes would be reproducible across models.

To profile and compare the cell types derived from each protocol, we first focused only on wild type (WT) isogenic control lines to avoid complication from potential *MAPT* mutation effects. We observed that Factory and HPC microglia were morphologically distinct from each other [Figure 1A-D, Figure S1A-C], with HPCs typically presenting as more spherical, while Factories had elongated, bipolar morphologies [Figure 1A]. Quantification of morphological features confirmed that while the average cell area across protocols was roughly equivalent [Figure 1B], HPCs were significantly rounder with increased width:length ratios and reduced perimeter lengths compared to the equivalent Factories [Figure 1B-D, Figure S1A-C].

**Figure 1.**
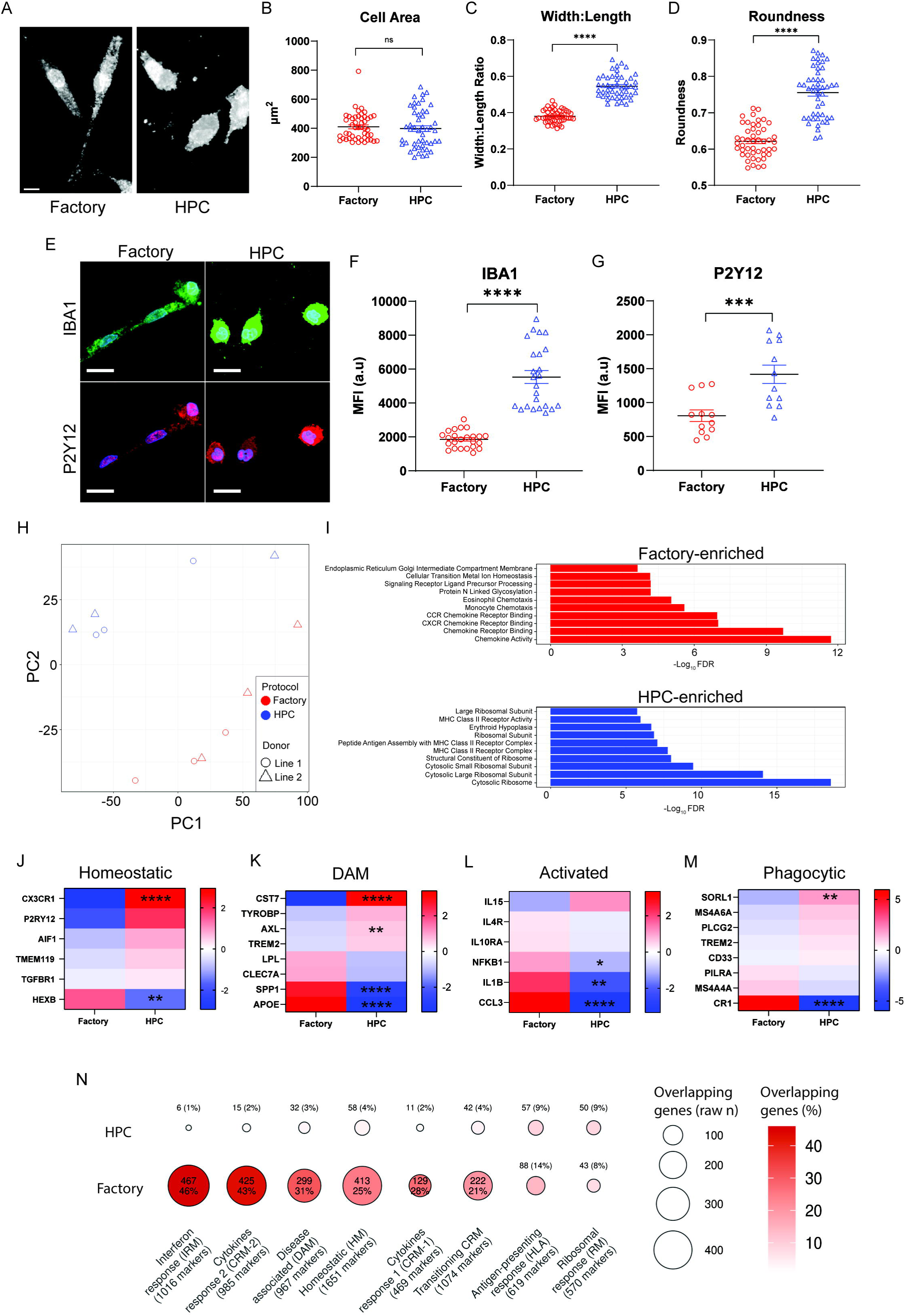
Factory and HPC protocols produce microglia in different functional states. **A.** Representative images of typical cellular morphologies following Factory or HPC microglia differentiation. Cells labelled with IBA1. Scale bar = 10µm. **B-D**. Quantification of microglia morphological features; ***B***. Cell area, ***C.*** Width:Length ratio and ***D.*** Roundness. n = 4 fields of view (FOV) from 4 independent wells per replicate. Data points represent mean value per FOV. Two-way ANOVA, Uncorrected Fisher’s LSD. **E.** Representative images of IBA1 and P2Y12 expression. Scale bar = 20µm. **F-G.** Quantification of ***F.*** IBA1 and ***G.*** P2Y12 mean fluorescence intensity (MFI) in arbitrary units (a.u). n = 4 FOV from 2 independent wells per replicate. Data points represent mean value per FOV. Two-way ANOVA, Uncorrected Fisher’s LSD. **H.** Principal components analysis of RNA-sequencing data. **I.** Top 10 Factory protocol-enriched (top, red) and HPC protocol-enriched (bottom, blue) gene ontology (GO) pathways. **J-K.** Z-score expression of ***J***. homeostatic, ***K.*** DAM, ***L.*** activated and ***M.*** phagocytic genes derived from RNA-sequencing data across protocols. **N.** Overlap between protocol-enriched differentially expressed genes and positive marker signature of each xenografted microglia state. Each row is a protocol gene set, and each column a microglial state. Dot size represents the number of overlapping genes, and the colour represents the percentage of that state’s markers recovered by protocol. All data derived from N = 2 independent donor iPSC lines, 3 independent differentiations per protocol. Error bars = SEM. *p < 0.05, **p < 0.01, ***p < 0.001, ****p < 0.0001, ns = not significant.

Cells derived from both protocols expressed microglial identity protein markers IBA1 and P2Y12 [Figure 1E-G], however HPCs exhibited significantly stronger fluorescence intensities for both compared to Factories [Figure 1F-G]. As expected from their morphological features, Factory and HPC microglia were transcriptionally distinct from each other [Figure 1H]. To further understand the differences between the two protocols, we then carried out differential gene expression and pathway enrichment analyses [Figure 1I, Figure S1D, Tables S1, S2]. Interestingly, we found that HPCs demonstrated upregulation of pathways related to ribosomal function and translation, as well as increased MHC Class II protein complexes, consistent with an antigen presenting-like state, and potentially associated with phagocytic responsiveness^36–39^ [Figure 1I, Table S2]. In contrast, Factories were enriched for pathways associated with cytokine and chemokine activity, including increased *Chemokine Receptor Binding* and *Monocyte Chemotaxis* pathways, compared to their HPC counterparts [Figure 1I, Table S2], signalling an increased inflammatory state, or increased potential for inflammation induction.

### Factory and HPC protocols result in microglia in different functional states

Given the apparent difference in microglial states between protocols, we then looked at the expression of specific marker genes [Figure 1J-M]. Similar to the imaging data, *P2RY12* expression was higher in HPCs compared to Factories, although this failed to reach statistical significance (log2FC = 1.98, p = 0.06). Similarly, there was a trend towards increased expression of other homeostatic identity markers in HPCs, including *TMEM119* (log2FC = 0.52), *TGFBR1* (log2FC = 0.26) and *AIF1* (log2FC = 0.82), although the fold change differences between these markers were minimal between protocols, and were not statistically significant [Figure 1J]. *CX3CR1* expression was significantly higher in HPCs (log2FC = 2.94, p < 0.001), possibly indicative of a more homeostatic state in these cells. Curiously, this was not consistent with the expression of microglia-specific marker *HEXB*, which was instead significantly more highly expressed in the Factories (log2FC = 1.56, p < 0.01) [Figure 1J]. While there was no consistent alteration in disease-associated microglia (DAM) gene expression, Factories had significantly higher expression of *APOE* (log2FC = 3.39, p < 0.001) and *SPP1* (log2FC = 2.75, p < 0.001) [Figure 1K]. On the other hand, *CST7* expression was significantly enriched in HPCs compared to the Factories (log2FC = 3.24, p < 0.001). Consistent with the pathway enrichment analysis, Factories also had significantly higher expression of several genes associated with microglial activation, namely *CCL3* (log2FC = 3.34), *IL1B* (log2FC = 2.21) and *NFKB1* (log2FC = 1.05), as well as interferon response gene *IFIT3* (log2FC = 3.92) [Figure 1L]. In contrast, phagocytosis gene *SORL1* was significantly higher in HPCs (log2FC = 1.90) [Figure 1M].

As our observed differences between microglia protocols were indicative of changes in state and function rather than overall microglial identity, we then compared their transcriptomic profiles to xenotransplanted iPSC-derived microglia^25^ in order to assess similarity to *in vivo* microglial states [Figure 1N, Figure S1E]. Unsurprisingly, both iPSC-microglia protocols showed very low similarity to *in vivo* microglia [Figure S1E]. However, when mapping the overlapping marker genes between *in vivo* microglia state with enriched genes for each protocol, we observed distinct enrichments [Figure 1N]. Specifically, Factory microglia showed a greater transcriptomic similarity to *in vivo* microglia, indicating these cells may be closer to an *in vivo* state than HPC microglia. Of each annotated state, Factory microglia were most strongly enriched for genes associated with interferon response and cytokine response [Figure 1N]. In contrast, HPC microglia showed the highest enrichment in antigen-presenting response and ribosomal response genes [Figure 1N], consistent with each protocol generating microglia representative of different functional states.

### The effect of *MAPT* mutation on microglia characteristics is protocol-dependent

After establishing the differences in HPC and Factory microglia in WT cells, we then examined whether *MAPT*-S305N impacted microglial identity, state, and gene expression across protocols. When examining cellular morphologies, *MAPT*-S305N had subtle but statistically significant effects [Figure 2A-C, Figure S2A-C], possibly reflecting cytoskeletal alterations as previously described in *MAPT*-IVS10+16 microglia^11^. *MAPT*-S305N led to significantly increased roundness across both protocols [Figure 2A-C], however the cause for this morphological change differed depending on protocol; HPC *MAPT*-S305N microglia were smaller and less complex [Figure S2A-C], whereas Factory *MAPT*-S305N microglia were wider than their isogenic controls [Figure S2A-C].

**Figure 2.**
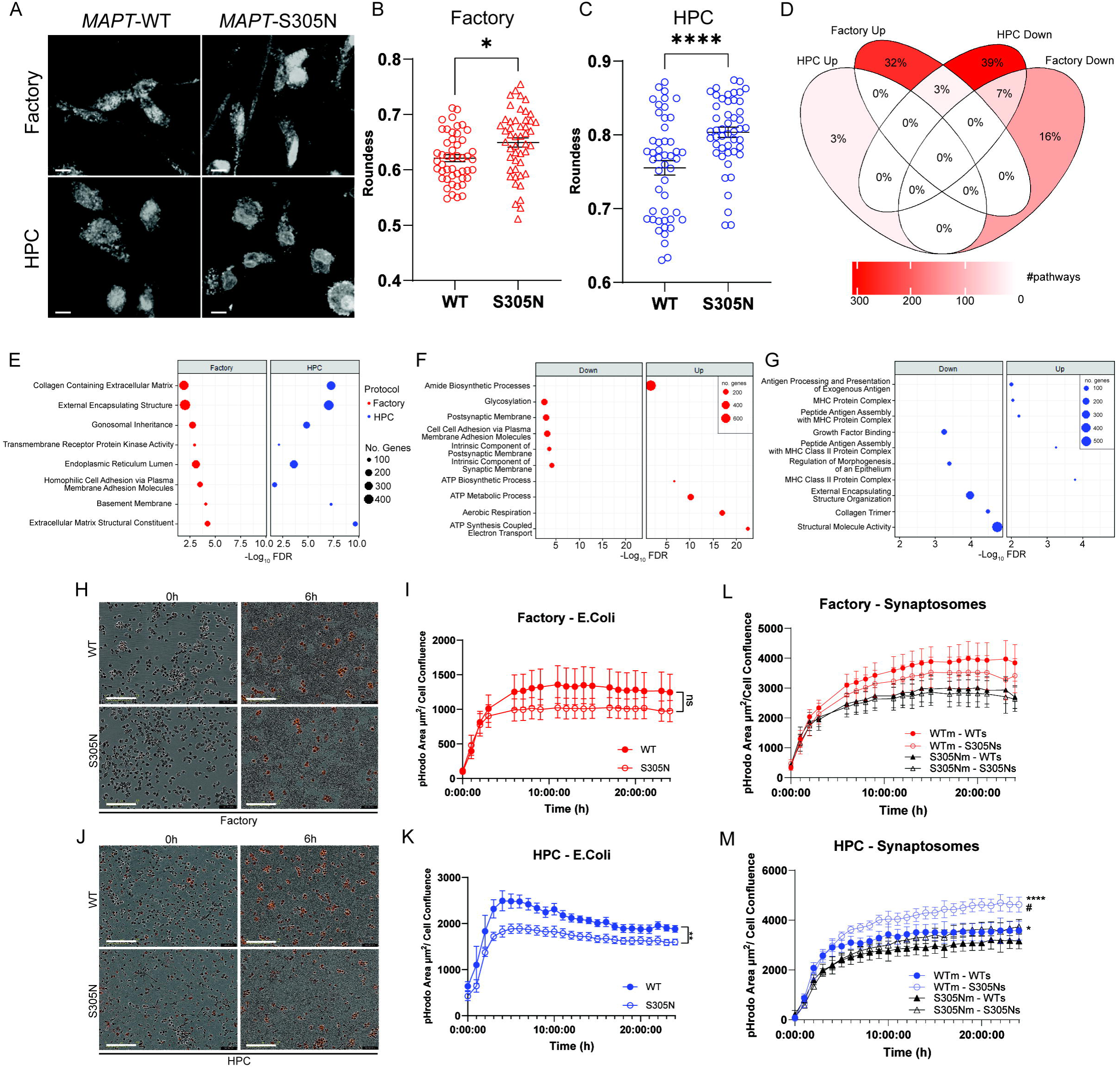
*MAPT* mutations in microglia alter extracellular matrix genes and impair phagocytosis. **A.** Representative images of typical cellular morphologies for each genotype following Factory or HPC microglia differentiation. Cells labelled with IBA1. Scale bar = 10µm. **B-C.** Quantification of cellular roundness between genotypes for ***B.*** Factory and ***C.*** HPC microglia. n = 4 fields of view (FOV) from 4 independent wells per replicate. Data points represent mean value per FOV. Two-way ANOVA, Uncorrected Fisher’s LSD. **D.** Overlap of the number of differentially expressed genes between *MAPT*-S305N microglia and isogenic controls across protocols. **E.** GO pathways significantly downregulated in *MAPT*-S305N microglia compared to isogenic controls in both protocols. **F-G.** Top 5 up- and down-regulated GO pathways in MAPT-S305N microglia following ***F.*** Factory and ***G.*** HPC protocols. **H**. Representative images of pHrodo red positive signal overlaid on brightfield images of microglia at baseline (0 hours) and 6 hours following E.Coli exposure in Factory microglia. **I.** Quantification of pHrodo positive area normalised to cell confluence over 24 hours following E.Coli exposure. Scale bar = 200µm. **J-K.** As ***H-I***, for HPC microglia. n = 2 independent wells per replicate. Mixed effects analysis. **L-M.** Quantification of pHrodo positive area normalised to cell confluence over 24 hours following treatment with either WT or *MAPT*-S305N organoid-derived synaptosomes in ***L.*** Factory and ***M.*** HPC microglia. In figure legends, ‘m’ refers to microglia genotype, and ‘s’ refers to synaptosome genotype. n = 2 independent wells per replicate. Multiple linear regression, Holm-Šídák’s multiple comparisons test. Asterisks denote statistical comparisons between synaptosome genotypes within the same microglia genotype. Hashes denote statistical comparisons between microglia genotype within the same synaptosome genotype. All data derived from N = 2 independent isogenic pairs of iPSC lines, 3 independent differentiations per protocol. Error bars = SEM. */#p < 0.05, **p < 0.01, ****p < 0.0001, ns = not significant.

Interestingly, the expression of homeostatic markers was altered by *MAPT*-S305N in HPCs only; fluorescence intensities of IBA1, P2Y12 and CD68 were all significantly increased in mutant cells compared to isogenic controls [Figure S2D-I]. However, this was not replicated in transcriptomic data, where there were no significant changes in the expression of homeostatic microglial genes nor any genes associated with microglial state due to the presence of *MAPT*-S305N in either protocol [Table S3].

After establishing that the expression of *MAPT*-S305N didn’t greatly impact microglial identity or state, we then explored the broader transcriptomic effect of mutation. After carrying out differential gene expression between *MAPT*-S305N and isogenic controls, we first looked at the overlap between significantly enriched pathways across protocols. Surprisingly, only 79 pathways (10% of all pathways) were significantly enriched in both HPC and Factory microglia, of which only 57 (7%) were altered in the same direction [Figure 2D]. These pathways were all downregulated, and converged on components of the extracellular matrix (ECM) [Figure 2E], suggesting that *MAPT*-S305N expression in microglia may contribute to ECM remodelling, with the potential to interfere with synaptogenesis^40,41^.

The majority of differentially regulated pathways due to *MAPT*-S305N were specific to the differentiation protocol employed [Figure 2D]. *MAPT*-S305N led to upregulation of metabolism and cellular respiration in Factories [Figure 2F, Table S4], similar to its previously reported impact in iPSC-neurons^17^. However, in microglia, upregulation of these processes has been proposed to be indicative of an anti-inflammatory or homeostatic phenotype^42^. In HPCs, *MAPT*-S305N amplified the differences between protocols, by upregulating antigen presentation-related pathways, primarily involving MHC and MHC class II [Figure 2G, Table S4], potentially indicating a shift towards a pro-inflammatory phenotype. Downregulated pathways in both protocols were consistent with the shared inhibitory effect of *MAPT*-S305N on ECM processes, involving collagen binding (HPC) and synaptic membrane components (Factories) [Figure 2E-G]

### *MAPT*-S305N causes phagocytic impairment

As phagocytosis is a crucial microglial function that can be altered by activation state^43^, we then assessed the phagocytic capacity of *MAPT*-S305N and isogenic control microglia derived from each protocol. We first challenged microglia with pHrodo-labelled E.Coli particles, and tracked their internalisation over 24 hours [Figure 2H-K, Figure S2J-K]. In HPC microglia, *MAPT*-S305N lead to significant impairment of E.Coli phagocytosis compared to isogenic controls [Figure 2J-K], in line with previous reports of impaired myelin phagocytosis in *MAPT*—IVS10+16 iPSC-microglia derived using the same protocol^11^. However, while there was a trend towards reduced phagocytic capacity in *MAPT*-S305N Factory microglia, this was not statistically significant [Figure 2H-I].

Microglia have been reported to contribute to the regulation of neuronal function by modulating synaptogenesis in health, and aberrantly engulfing synapses in disease^44–46^. Indeed, our transcriptomic data was indicative of impaired regulation of synaptic support in *MAPT*-S305N microglia [Figure 2F]. In order to determine whether *MAPT*-S305N microglia differentially responded to synapses in a tauopathy context, we isolated and labelled synaptosomes from 6-month-old *MAPT*-S305N and isogenic control cortical organoids [Figure S2L]. As with E.Coli, we observed impaired synaptosome internalisation in *MAPT*-S305N microglia across both protocols, although this only reached statistical significance in HPC microglia [Figure 2L-M]. Interestingly, there was only an interaction between synaptosomes containing pathogenic tau and microglia genotype in HPCs [Figure 2M]. Specifically, control microglia internalised synaptosomes containing pathogenic tau at a significantly faster rate than synaptosomes containing with WT tau [Figure 2M], indicating that microglia were able to detect the presence of pathogenic proteins in synapses and could respond effectively. In contrast, *MAPT*-S305N microglia were less able to make this distinction [Figure 2M], and therefore may be less likely to effectively clear pathogenic proteins or dysfunctional synapses. This effect was not present in Factories, which did not differentially respond to synaptosome genotype [Figure 2L]. Overall, while microglia derived from either protocol were capable of phagocytosis and broadly replicated impairment due to *MAPT*-S305N, we noted that HPCs consistently showed a greater capacity for internalisation, with pHrodo fluorescence intensities being ∼1.5 – 2x fold higher compared to Factories [Figure 2I-K, L-M], consistent with their increased expression of phagocytosis genes [Figure 1M]. However, as the expression of phagocytosis genes were no different between *MAPT*-S305N microglia and isogenic controls in either protocol [Table S3], the effect of pathogenic tau on phagocytosis is likely post-transcriptional.

### Factory microglia have a greater response to inflammatory stimuli

Neuroinflammation is a key pathogenic hallmark of numerous neurodegenerative diseases, including FTD-tau^6,47^. Furthermore, our prior work in iPSC-astrocytes suggests that the *MAPT*-S305N mutation induces a pro-inflammatory state under basal conditions^17^. We therefore sought to determine whether *MAPT*-S305N also alters inflammatory processes in iPSC-microglia. In order to do this, we challenged microglia with either a pro-inflammatory (lipopolysaccharide (LPS)) or anti-inflammatory (IL4) stimulus, and examined cytokine/chemokine release and transcriptomic responses.

When examining the transcriptomic data, principal components analysis revealed that LPS/IL4 treatment had a clear and opposing transcriptomic effect in Factories [Figure 3A], but not in HPCs, where donor line explained separation along PC2 [Figure 3B]. This was consistent with our earlier observation of gene expression differences between protocols, in which Factories had increased expression of cytokine and chemokine receptor genes compared to HPCs [Figure 1I], and thereby may be more primed to respond to stimulation. Consistently, both WT and *MAPT*-S305N HPCs showed fewer differentially expressed genes following stimulation compared to their Factory counterparts, with LPS surprisingly being unable to induce any significant transcriptional change in HPCs [Figure S3A-B].

**Figure 3.**
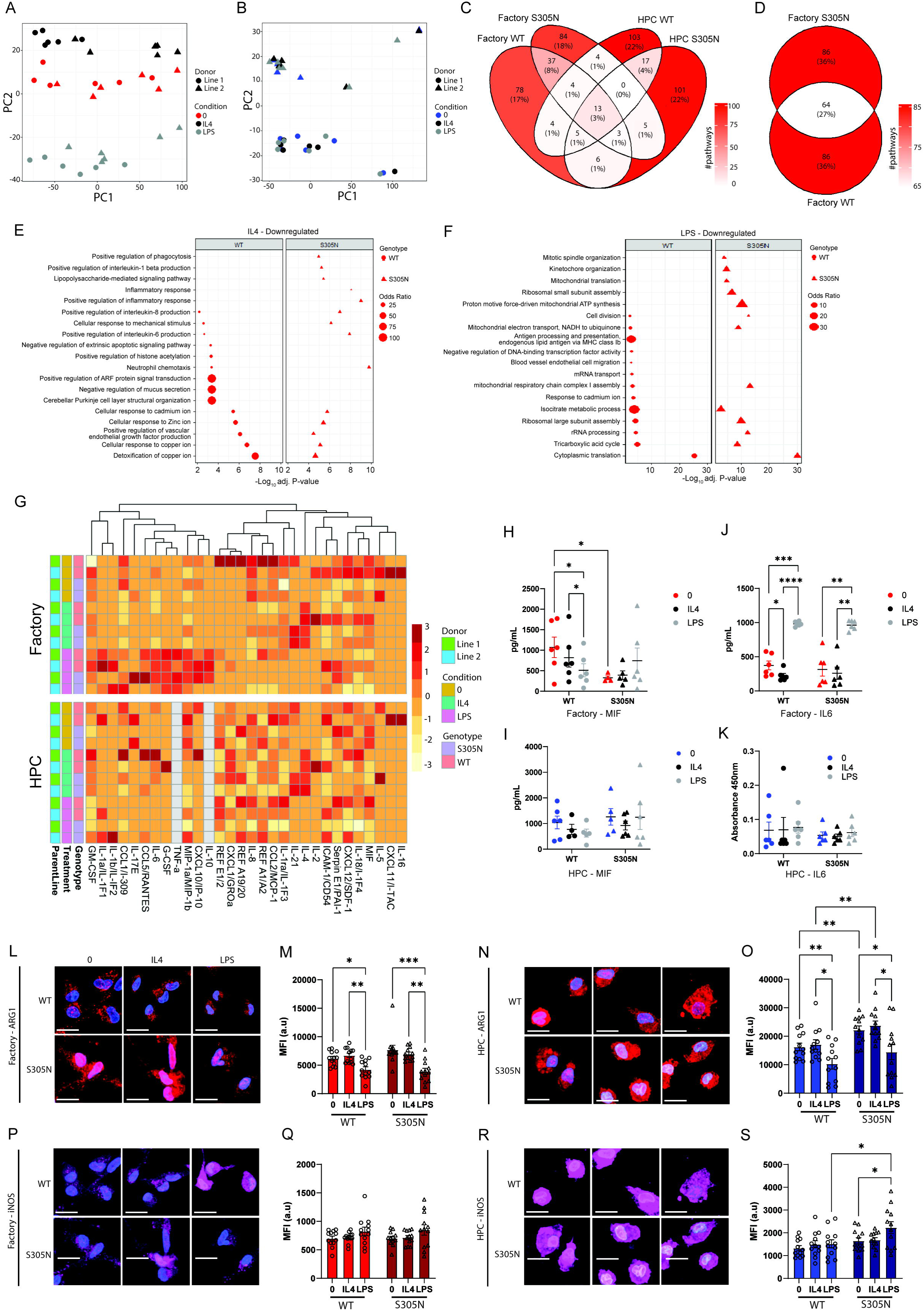
Factory microglia better respond to inflammatory stimuli, but *MAPT*-S305N has a minimal effect. **A-B.** PCA analysis of RNA-sequencing data in ***A.*** Factory and ***B.*** HPC microglia, following exposure to either LPS or IL4. **C.** Overlap of differentially expressed genes following IL4 stimulation compared to untreated cells across each genotype and protocol. **D.** Overlap of differentially expressed genes in WT and *MAPT*-S305N Factory microglia following LPS stimulation compared to untreated cells. **E-F.** Significantly downregulated GO pathways in WT and MAPT-S305N Factory microglia following ***E.*** IL4 and ***F***. LPS stimulation. **G.** Relative cytokine detection following cytokine array analysis in Factory (top) and HPC (bottom) microglia conditioned media. Detection values were scaled by column, and cytokines were arranged by unsupervised hierarchical clustering. Data derived from one microglia differentiation per cell line per protocol. **H-K.** Quantification of ***H-I.*** MIF and ***J-K***. IL6 under different stimulation conditions across genotypes by ELISA in ***H,J*** Factory and ***I,K*** HPC microglia conditioned media. Mixed effects analysis, Tukey’s multiple comparisons test. **L-O.** Representative images ***(L, N)*** and quantification ***(M, O)*** of ARG1 MFI in ***L-M***. Factory and ***N-O*** HPC microglia following inflammatory stimuli. **P-S.** Representative images ***(P, R)*** and quantification ***(Q, S)*** of iNOS MFI in ***P-Q***. Factory and ***R-S*** HPC microglia following inflammatory stimuli. Scale bar = 20µm Unless otherwise stated, A- Error bars = SEM. */p < 0.05, **p < 0.01, ***p < 0.001, ns = not significant.

We next evaluated whether *MAPT*-S305N significantly altered microglial responses to inflammatory stimuli in either protocol. Overall, there were very few significantly enriched pathways in common between genotypes or across protocols following IL4 stimulation [Figure 3C], although the proportion of common pathways across genotypes in Factories was higher than that in HPCs (13% vs 8%, respectively). There was greater concordance across genotypes following LPS stimulation, with 27% of all significantly enriched pathways shared between *MAPT*-S305N and isogenic control Factories [Figure 3D].

Although there was not a large proportion of enriched pathway overlap between genotypes, following IL4 exposure the top pathways altered in both *MAPT*-S305N and isogenic control Factories were similar; both showed significant downregulation of pathways associated with ion responses and interleukin production, and upregulation of antigen response and chemotaxis pathways [Figure 3E, Figure S3C]. However, the reduction in inflammatory signalling occurred to a greater extent in *MAPT*-S305N Factories, which also had significant downregulation of additional pathways including *lipopolysaccharide-mediated signaling pathway, inflammatory response,* and *positive regulation of inflammatory response,* which were not significantly enriched in WT cells [Figure 3E, Table S5], perhaps indicating an amplified anti-inflammatory response to IL4 in mutant cells. IL4 treatment of HPCs of either genotype resulted in vastly different transcriptomic changes compared to Factories; downregulated pathways were not aligned with microglial-specific functions and likely reflect spurious results [Figure S3D]. Indeed, almost all significantly enriched pathways were due to enrichment of a single gene within small annotation sets. While upregulated pathway enrichments were more robust, they were indicative of upregulated inflammatory responses rather than the predicted anti-inflammatory effects following IL4 exposure [Figure S3E]. The effect of *MAPT*-S305N on the IL4 response in HPCs was unsurprisingly minimal.

As suggested by the PCA analyses [Figure 3A-B], there were no significantly differentially expressed genes in HPCs following LPS exposure for either genotype [Figure S3B]. As such, it was not possible to conduct pathway enrichment. In Factories, the top upregulated pathway in both *MAPT*-S305N and isogenic control cells following LPS exposure was reassuringly *cellular response to lipopolysaccharide* [Figure S3F, Table S5]. Both genotypes responded to LPS with upregulation of pathways associated with inflammation [Figure S3F], while supressing translation [Figure 3F]. Interestingly, *MAPT*-S305N Factories appeared to be more prone to proliferation impairment resulting from LPS exposure compared to isogenic controls, with downregulation of *mitotic spindle organization* and *kinetochore organisation*, as well as a greater enrichment of genes associated with *cell division* compared to isogenic controls [Figure 3F, Table S4], indicating that *MAPT* mutations may have subtle effects on microglial responses to stimulation.

### *MAPT*-S305N has minimal impact on microglial cytokine responses to inflammatory stimuli

To further examine the differential effects of both microglia protocol and *MAPT* mutation following stimulation, cytokine and chemokine release into the media was assessed. In line with the transcriptomic data, we observed different basal cytokine profiles across protocols [Figure 3G-K]. Under basal conditions, *MAPT*-S305N Factories excreted reduced levels of MIF, IL-18, CXCL12 and Serpin E1 compared to their isogenic controls [Figure 3G], potentially indicating a suppression of pro-inflammatory processes in the presence of the mutation. In contrast, this pattern was not observed in HPC-microglia. Indeed, basal levels of MIF were moderately higher in *MAPT*-S305N HPC-microglia compared to isogenic controls [Figure 3G]. We therefore chose to quantify MIF secretion by ELISA, which was consistent with the cytokine array observations [Figure 3H-I].

As observed in the transcriptomic data, there was no apparent effect of LPS stimulation in HPCs, whereas Factories were able to mount a robust response with excretion of expected factors including TNFα and IL6 [Figure 3G]. Quantification of IL6 by ELISA confirmed these findings, with a significant upregulation in response to LPS in Factories [Figure 3J], whereas IL6 detection in HPCs remained below the sensitivity of the assay, despite repetition with double the input [Figure 3K]. However, we did not observe any consistent genotype-specific effects in the LPS cytokine response.

Finally, there was no consistent effect of IL4 stimulation on cytokine release, with the exception of IL4 detection itself, and IL21 [Figure 3G]. Interestingly, we observed increased IL21 release only in *MAPT*-S305N neurons following IL4 stimulation across both microglia protocols. This effect was not replicated when assessed by ELISA as detection was consistently below the sensitivity of the assay, although there was a trend towards increased absorbance values in Factory *MAPT*-S305N microglia compared to isogenic controls following IL4 stimulation [Figure S3G-H].

Lastly, to verify whether *MAPT*-S305N microglia differed in their response to inflammatory stimuli, we visualised levels of the anti-inflammatory marker ARG1 and pro-inflammatory marker iNOS. ARG1 expression in HPCs was roughly double that in Factory microglia [Figure 3L-O], consistent with them being closer to a state of efferocytosis than inflammation. While ARG1 levels did not appear to be responsive to IL4, they did significantly drop across all genotypes and protocols in response to LPS stimulation [Figure 3L-O]. Furthermore, *MAPT*-S305N HPCs had significantly higher levels of ARG1 in basal and IL4 conditions in comparison to their isogenic controls, consistent with suppressed inflammatory processes in mutant cells, although this difference was absent following LPS [Figure 3N-O]. iNOS levels were less responsive to stimulation than expected, with only minor trends towards an LPS- induced increase in Factories [Figure 3P-Q]. There was a small yet significant increase of iNOS in *MAPT*-S305N HPCs in response to LPS, which was also significantly higher in comparison to their isogenic controls [Figure 3R-S]. This indicated that despite the lack of transcriptomic or cytokine release responses, HPCs are at least capable of minimally responding to both IL4 and LPS stimulation [Figure 3N-O, R-S].

### *MAPT* genotype influences microglial response to *MAPT* mutation neurons

We next carried out mixed genotype neuron-microglia co-cultures in order to assess potential non-cell autonomous effects of *MAPT*-S305N in microglia and neurons. For these experiments, we focused our attention only on Factories, as in our hands and for these particular iPSC lines, we found this protocol to be more reliable and scalable for microglia production. Neurons were generated using directed differentiation^27^, resulting in production of a mixed pool of neurons and astrocytes. Addition of microglia therefore resulted in tri-cultures [Figure 4A-B]. Mixed genotype tri-cultures underwent single cell sequencing, and both neuronal and microglial transcriptomic changes were assessed in the presence of cells of different *MAPT* genotypes [Figure 4A].

**Figure 4.**
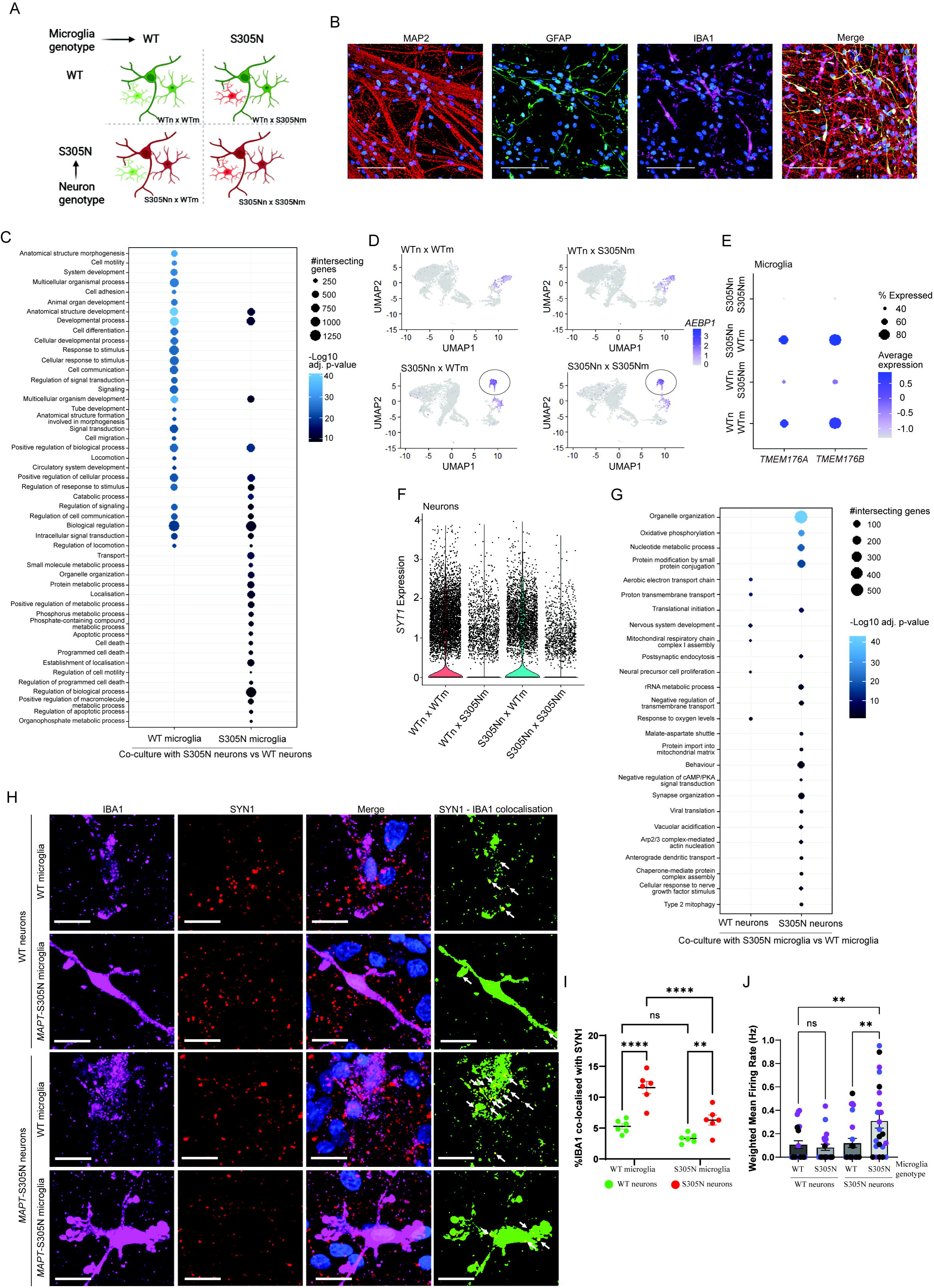
*MAPT-S305N* microglia differentially respond to pathogenic neurons and influence synaptic function. **A.** Schematic of mixed genotype neuron-microglia co-culture design. **B.** Representative images of tri-cultures, labelled with MAP2 (neurons), GFAP (astrocytes) and IBA1 (microglia), each with DAPI nuclear stain (blue). Scale bar = 100µm. **C.** Upregulated GO pathways in microglia when co-cultured with either MAPT-S305N or WT neurons. **D.** UMAP projection of microglia under mixed genotype co-culture conditions. Circle in bottom two panels indicates microglia population unique to MAPT-S305N neuron co-culture. **E.** TMEM176A and TMEM176B expression in microglia under different mixed genotype co-culture conditions. **F.** Expression of SYT1 in neurons under different mixed genotype co-culture conditions. **G.** Upregulated GO pathways in neurons when co-cultured with either MAPT-S305N or WT microglia. Differential gene expression determined using Wilcoxon rank-sum test, Bonferroni correction. Sequencing data derived from N = 2 independent isogenic pairs of iPSC lines from one round of differentiation (n = 24,197 microglia, 34,962 neurons). **H.** Representative images of IBA1 and Synapsin I (SYN1) co-localisation under different co-culture conditions. White arrows in the right-hand panels indicate points of co-localisation between IBA1 and Synapsin I. Scale bar = 20 µm. I. Percentage of IBA1 area co-localized with Synapsin I under different genotype co-culture conditions. N = 1 isogenic pair of iPSC lines from three independent rounds of differentiation, with 2 replicate wells analysed per batch. Two-way ANOVA, uncorrected Fisher’s LSD multiple comparisons test. **J.** Weighted mean firing rate of mixed genotype co-cultures determined by multi-electrode array (MEA). Different coloured datapoints represent independent differentiations. N = 2 isogenic pairs, three independent rounds of differentiation, 8 replicate wells per batch. One-way ANOVA, Holm-Šídák’s multiple comparisons test. Error bars = SEM. *p < 0.05, **p < 0.01, ***p < 0.001, ns = not significant.

When examining microglia co-cultured with either control or *MAPT*-S305N neurons, we found that both control and *MAPT*-S305N microglia responded to *MAPT* mutation neurons by upregulating pathways associated with regulation of cellular communication, as well as migration and locomotion, in comparison to when cultured with control neurons [Figure 4C]. Interestingly, co-culture with *MAPT*-S305N neurons resulted in a unique cluster of microglia that was not present in control neuron co-cultures, despite microglia being derived from the same differentiations and collections at the same time for both co-culture conditions [Figure 4D]. The top three marker genes for this cluster were *AEBP1*, *ITGB4* and *TPPP3*, suggesting that *MAPT*-S305N neurons induced a population of pro-inflammatory, activated microglia regardless of microglia genotype [Figure 4D, Figure S4D-E]. Pathway enrichment of marker genes in this cluster were indicative of functions associated with extracellular matrix and cytoskeleton, consistent with a change in microglial function [Figure S4F].

However, microglia genotype modified the way in which they responded to neuronal genotype. While control microglia responded to mutant neurons by upregulation of pathways related to stimulus responses, *MAPT*-S305N microglia failed to do so [Figure 4C]. For example, *MAPT*-S305N and control microglia showed differential expression of *TMEM176B* and *TMEM176A* expression when cultured with *MAPT*-S305N neurons, with both being suppressed in *MAPT*-S305N microglia, while they were induced in control microglia [Figure 4E]. Imaging of co-cultures confirmed significantly suppressed TMEM176B expression in *MAPT*-S305N microglia compared to isogenic controls when cultured with WT neurons [Figure S4G]. In contrast to the transcriptomic data, *MAPT*-S305N neurons caused a significant reduction in TMEM176B detection by immunofluorescence in WT microglia, and had no impact on *MAPT*-S305N microglia [Figure S4G]. While discordant with the gene expression data, these findings still supported impaired TMEM176B expression in *MAPT* mutation microglia and differential responsiveness to neurons. Indeed, *MAPT*-S305N microglia instead responded to *MAPT*-S305N neurons with upregulation of pathways associated with a switch in metabolic processes and cell death [Figure 4C], indicating that *MAPT* genotype influences the way in which microglia respond to a pathogenic environment.

### Neuronal transcriptomes are influenced by microglia *MAPT* genotype

We next examined whether microglial *MAPT* genotype was able to influence neuronal transcriptomes and function. As expected, microglia genotype did not have a large impact on neuronal transcriptomes, with only 95 genes being differentially expressed in control neurons when cultured with *MAPT*-S305N compared to control microglia. However, *MAPT*-S305N neurons were much more susceptible to the influence of microglia genotype, with 2199 genes significantly differentially expressed in response to *MAPT*-S305N microglia.

In both control and *MAPT*-S305N neurons, the top differentially expressed gene was *SYT1*, which was downregulated when neurons were cultured with *MAPT*-S305N compared to control microglia, indicative of a potential effect of microglial genotype on neuronal activity [Figure 4F]. Additionally, when cultured with control neurons, microglia genotype influenced neuronal metabolism; *MAPT*-S305N microglia caused upregulation of several mitochondrial electron transport chain pathways in neurons compared to when they were co-cultured with control microglia [Figure 4G]. Oxidative phosphorylation was also upregulated in *MAPT*-S305N neurons when co-cultured with *MAPT*-S305N microglia, as well as additional pathways related to neuronal communication and function, such as *synapse organisation*, *postsynaptic endocytosis* and *anterograde dendritic transport* [Figure 4G], suggesting that microglial *MAPT* genotype may more greatly influence neuronal signalling when in a pathogenic context.

### *MAPT*-S305N mutation microglia contribute to neuronal hyperexcitability by impaired synaptic regulation

Combining our initial transcriptomic data indicating the *MAPT*-S305N mutation may interfere with synapse regulation via impairment of the extracellular matrix [Figure 2E], and phagocytosis data indicating impaired synaptic engulfment [Figure 3L], we therefore hypothesised that *MAPT*-S305N microglia may fail to properly regulate neuronal synaptogenesis by having a reduced capacity to respond to and engulf aberrantly formed synapses, or synapses containing pathogenic tau. To test this, we carried out immunofluorescence labelling of mixed genotype co-cultures with synaptic marker Synapsin I and examined its co-localisation with microglial marker IBA1 [Figure 4H-I]. Consistent with our hypothesis, control microglia showed a significantly increased colocalization with Synapsin I puncta when co-cultured with *MAPT*-S305N neurons compared to when they were cultured with control neurons [Figure 4H-I], indicative of increased synaptic engulfment in response to the presence of pathogenic tau. In contrast, while *MAPT*-S305N microglia were capable of responding to the presence of *MAPT*-S305N neurons, they showed significantly less Synapsin I co-localisation compared to WT microglia [Figure 4H-I], consistent with a phagocytic impairment that results in a reduced ability to regulate aberrant synaptic activity.

Subsequently, we measured spontaneous neuronal activity in co-cultures by multi-electrode array [Figure 4J]. *MAPT*-S305N neurons cultured with *MAPT*-S305N microglia exhibited significantly increased spontaneous activity compared to control neurons cultured with control microglia, consistent with hyperexcitability phenotypes observed in *MAPT* mutation iPSC-neurons by our group and others^17,48–50^, and an inability for *MAPT* mutation microglia to appropriately regulate synaptic function. However, while *MAPT*-S305N microglia had no impact on control neuronal activity, co-culture of *MAPT*-S305N neurons with control microglia was able to restore spontaneous neuronal activity to control levels [Figure 4J], consistent with our observations that these microglia displayed improved clearance of synapses containing pathogenic tau protein.

## DISCUSSION

We have carried out transcriptomic and functional profiling of the impact of the *MAPT*-S305N mutation in iPSC-microglia derived from two different protocols in parallel. As a result, we have provided the first reported direct comparison of two of the most popular and widely used microglia differentiation approaches, Haenseler *et al*^16^ (Factories) and McQuade *et a*l^15^ (HPC). This comparison resulted in the observation that each protocol generates microglia in different functional states, and importantly, these basal states defined the observable pathogenic phenotypes that resulted from *MAPT* mutations. Interestingly, the effect of *MAPT*-S305N across both protocols was consistent with a suppression of microglial reactivity, evidenced by impaired phagocytosis, reduced cytokine release and diminished regulation of neuronal synaptic function. As such, we propose that *MAPT* mutation microglia likely contribute to disease pathogenesis by their inability to effectively respond to pathogenic environments and their failure to protect vulnerable neurons.

While we did not directly examine why each differentiation protocol should result in different microglial states, the use of different media components during microglia differentiation from the initial progenitor stages likely contribute. The Factory protocol was initially developed for the production of yolk-sac derived macrophages^51^, with subsequent iterations leveraging the protocol’s generation of *MYB*-independent myeloid progenitors to direct cells towards a microglial fate^16,52^. Similarly, HPC microglia recapitulate differentiation to primitive hematopoietic cells resulting from yolk-sac erythromyeloid progenitors, prior to microglial specification^16,53^, thereby reflecting similar initial progenitor pools and microglial ontologies. Regardless, the protocols significantly diverge during microglia maturation, with inclusion of multiple factors with potential to impact microglial state. For example, insulin has been identified as a regulator of microglia metabolism, with insulin resistance associated with impaired amyloid β (Aβ) phagocytic uptake^54^, whereas insulin replacement can enhance this function and suppresses microglial activation in models of AD^55,56^. As such, the inclusion of insulin in the HPC protocol may be contributing to their less reactive, pro-phagocytic phenotype. In contrast, the Factory protocol includes low levels of GM-CSF in the microglia maturation media; while the inclusion of this cytokine has been reported to support microglial adherence and proliferation^52^, it has also been reported to promote a pro-inflammatory or primed microglial state^57,58^, which may contribute to the enhanced inflammatory reactivity we observe in these cells. However, it is important to note that as we did not deviate from published Factory or HPC protocols or test different media conditions, the contribution of these components to the observed divergent microglial states is not definitive. Regardless, the functional profile of iPSC-microglia should be an important consideration when selecting which protocol to adopt for planned investigations, depending on the phenotype of interest.

When comparing the impact of the pathological *MAPT*-S305N variant, we found very few similarities across protocols, with observable phenotypes typically in alignment with the microglial starting state. For example, while both protocols supported a defect in phagocytosis in *MAPT*-S305N microglia, this only reached statistical significance in HPC microglia. Similarly, we were able to detect a deficit in the recognition of pathogenic tau-containing synaptosomes due to *MAPT*-S305N only in the HPC microglia, indicating that cells already in a pro-phagocytic state are more likely to show subtle functional differences undetectable in other, less specialized cells. These findings replicate recent reports from *MAPT* IVS10+16 iPSC-derived microglia, which showed impaired phagocytic engulfment of myelin and recombinant tau protein^11^, indicating that this impairment is broadly applicable to a range of different neurodegenerative disease-relevant substrates. Given that there were minimal transcriptomic changes due to *MAPT* mutation in microglia derived from either protocol, none of which were directly relevant to phagocytosis, phagocytic impairment is most likely to occur post-transcriptionally. Microtubule destabilization and disorganization due to the presence of pathogenic tau is a compelling explanation, which would alter the ability for cells to effectively engulf and transport cargo^59^. This hypothesis is supported by our observation of altered microglial morphologies in *MAPT*-S305N cells, and by disorganized tubulin cytoskeletons in *MAPT*-IVS10+16 microglia observed by others^11^. However, *MAPT-*S305N has been shown to accelerate phagocytic activity in iPSC-astrocytes^17^, therefore altered interactions between pathogenic tau and microtubules are unlikely to be the sole cause of microglial phagocytic impairment.

Microglial microtubule organization has also been implicated in microglial reactivity and activation^60,61^. Interestingly, we observed a trend towards reduced cytokine and chemokine release in *MAPT*-S305N microglia under basal conditions, supporting a suppression of microglial reactivity. However, mutant microglia were still capable of responding to LPS stimulation, indicating that *MAPT*-S305N does not broadly impair responsiveness to strong inflammatory stimuli. Yet, co-culture with *MAPT* mutation neurons revealed alterations in microglial responsiveness to less aggressive pathogenic stimuli; while a subset of *MAPT-*S305N microglia showed capacity to transition to a pro-inflammatory activated state, the majority of microglia failed to appropriately respond to the presence of pathogenic tau and instead upregulated pathways related to cell death, indicating *MAPT* mutation microglia may be less likely to provide necessary neuroprotection under stress. The lack of an effect of *MAPT*-S305N microglia on WT neurons indicates that *MAPT* microglial genotype most likely influences disease processes only under pathogenic conditions when neurons are already vulnerable, rather than independently driving pathogenesis.

Our observations of reduced reactivity and phagocytic capability are seemingly at odds with current understanding of microglial roles in neurodegenerative disease, which describe chronic over-activation of neuroinflammatory processes and aberrant engulfment of synapses and neurons, subsequently contributing to neuronal dysfunction and loss^10,12,62–65^. Instead, we propose a model where *MAPT* mutations lead to microglial hypofunction, evidenced by an impaired ability to phagocytose synaptosomes containing pathogenic tau, a suppressed basal inflammatory state, and a reduced ability to mount an appropriate response when cultured with pathogenic neurons. This hypoactivation subsequently results in an inability to effectively regulate synaptic development and activity or protect neurons from pathogenic tau, thereby supporting neuronal hyperexcitability and conferring additional damage and stress to already vulnerable neurons. However, given that neuroinflammation is a characteristic feature of primary tauopathies and correlates with disease progression^6^, we hypothesise that our model reflects early pathogenic mechanisms that occur while neurons are not yet undergoing active synaptic loss and cell death. Indeed, *MAPT*-S305N microglia were capable of responding to strong neuroinflammatory stimuli similarly to control microglia. As such, microglial contributions to disease may reflect a two-stage process; an initial stage where hypofunctional microglia enhance later neuronal vulnerability through ineffective protection and synaptic regulation, followed by a second pathogenic stage where microglia mount a robust, sustained and toxic inflammatory response to neuronal death and dysfunction. Such temporal alterations in microglia state and reactivity correlating with pathological severity have been observed in human post-mortem brain tissue^9^. Whether this hypothesised switch in state could be observed *in vitro* likely requires longer cultures subjected to more chronic stressors than were conducted here

The question remains regarding how or why *MAPT* genotype should influence microglia function at all, given its very low expression in these cells. Other than the potential influence on microtubule organization, another intriguing possibility is the concept of tau “fatigue,” by which tau-exposed microglia subsequently become hypofunctional. For example, primary rodent microglia exposed to pathogenic tau became hypophagocytic^12^. Interestingly, peripheral monocytes derived from *MAPT* mutation carriers have been found to express reduced *TMEM176A* and *TMEM176B*^66^, which we also observed in our *MAPT*-S305N mutation microglia, and is consistent with suppressed inflammatory signalling in myeloid lineage cells in response to encountering pathogenic tau. Given that we observe these effects in monoculture microglia in the absence of any other stimuli, it is possible that even very low levels of endogenously expressed pathogenic tau are sufficient to predispose microglia to hypoactivity. How tau exerts such an effect remains to be investigated, however given that fibrillar tau appears to induce a toxic neuroinflammatory response^7,8^, any suppressive effect of tau exposure is most likely to occur prior to aggregation events.

Overall, we conclude that neither tested differentiation protocol is superior to the other with regards to the generation of microglia-like cells, however experimental requirements ought to guide selection of the most appropriate approach to use, depending on phenotypes of interest. Despite these protocol differences, this work has revealed novel insights regarding the cell autonomous and non-cell autonomous effects of *MAPT* mutations in microglia. Specifically, we find that endogenous expression of pathogenic tau induces microglial hypoactivity, such that cells fail to adequately respond to pathogenic stimuli or regulate aberrant synaptic activity, therefore contributing to early stages of disease pathogenesis and neuronal vulnerability.

## Supplementary Tables

*Table S1*. Significantly differentially expressed genes in HPC vs Factory control microglia

*Table S2.* Top 10 GSEA Pathway enrichments in HPC and Factory control microglia

*Table S3*. Differentially expressed genes in *MAPT*-S305N vs isogenic control microglia derived from each protocol

*Table S4.* Significantly enriched GO pathways in *MAPT*-S305N microglia compared to isogenic controls for each protocol

*Table S5.* Significant GO:BP Pathway enrichments following inflammatory stimuli across genotypes and microglia protocols

## Supplementary Figures

*Figure S1.* Factory and HPC protocols result in microglia in different functional states, and are both transcriptomically distinct from primary adult human microglia. Related to Figure 1.

*Figure S2. MAPT*-S305N causes mild morphological changes in microglia derived from both protocols. Related to Figure 2.

*Figure S3.* Factory and HPC protocol microglia have distinct responses to inflammatory stimuli. Related to Figure 3.

*Figure S4.* A population of microglia from both genotypes are capable of responding to *MAPT* mutation neurons, but *MAPT*-S305N microglia show impaired TMEM176B expression. Related to Figure 4.

## Data Availability

All transcriptomic data will be made available via the Gene Expression Omnibus upon final publication, and outputs of the analyses will be available on request. All code used in this study has been previously published, and pipelines are described in the Methods. Any other additional data or information reported in this publication is available from the lead contact upon request.

## Acknowledgements

We are grateful to the Tau Consortium and Rainwater Charitable Foundation for supporting this project. We thank the research subjects and their families for their generous participation, including in ARTFL (U54NS092089) and LEFFTDS (U01AG045390) that has made the generation of the *MAPT*-S305N iPSC lines possible. This work was supported by funding from the Rainwater Charitable Foundation (KRB, LKK, SH), Medical Research Scotland (CL), and the UK Dementia Research Institute (UKDRI-4211) through UK DRI Ltd, principally funded by the UK Medical Research Council (KRB, OD). We thank Meryam Benniazza and Viktoria Major at the University of Edinburgh Single Cell Facility for their assistance and input, as well as the High Content Screening Facility and Clinical Research Facility at the University of Edinburgh for their services and support. Figure 4A was created with BioRender.

## Author Contributions

Conceptualization: KRB

Methodology: DV, LKK, CL, TW, SH, JCW, OD, KRB

Validation: LKK, CL, KRB

Formal Analysis: DV, TW, JCW, OD, KRB Investigation: DV, LKK, CL, TW, JCW, OD, KRB Resources: RM, KRB

Data Curation: DV, OD, KRB Writing – Original Draft: KRB

Writing – Review and Editing: DV, LKK, CL, TW, SH, JCW, OD, RM, KRB Visualization: DV, CL, TW, JCW, KRB

Supervision: RM, KRB Funding Acquisition: KRB

