## Supplemental Figures for "Cross-protocol comparison of iPSC-microglia reveals hypofunction contributes to neuronal vulnerability and synaptic alterations in the *MAPT*-S305N model of frontotemporal dementia"

### Supplementary Information

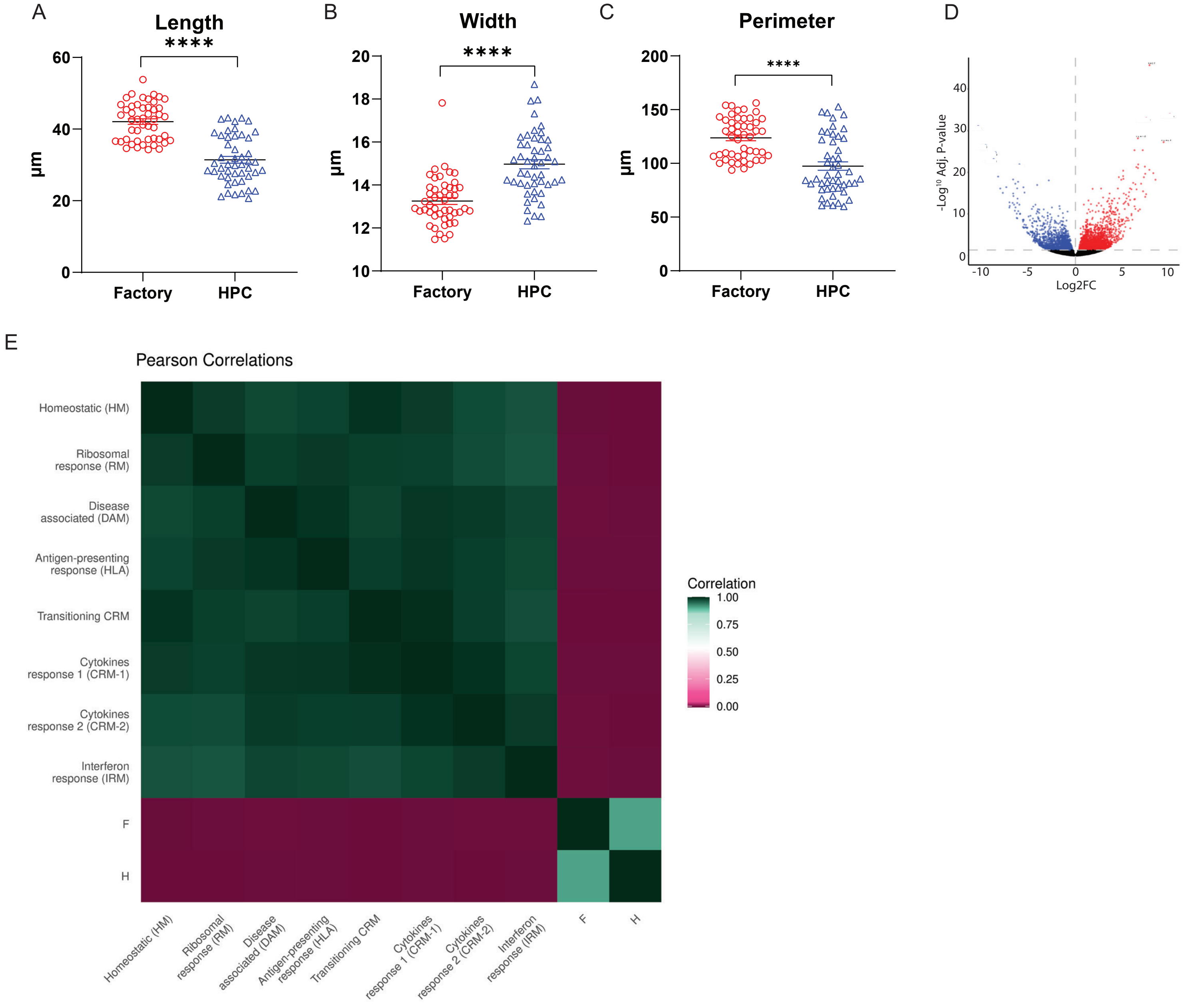


**Figure S1. Factory and HPC protocols result in microglia in diffrerent functional states, and are both transcriptomically distinct from primary adult human microglia. Related to Figure 1.**

**A - C.** Quantification of morpholigical features of Factory and HPC microglia. n = 4 fields of view (FOV) from 4 independent wells per replicate. Data points represent mean value per FOV. Two-way ANOVA, Uncorrected Fisher's LSD.

**D**. Extent of differential gene expression between WT HPC and Factory microglia derived from the same iPSC lines.

**E**. Pairwise Pearson correlation coefficients computed across averaged gene expression profiles, restricted to the 17,984 genes shared by all datasets. Colour encodes the correlation coefficient (green, high; purple, low). F = Factory microglia, H = HPC microglia

All data derived from N = 2 independent donor iPSC lines, 3 independent differentiations per protocol. Error bars = SEM. *p < 0.05, **p < 0.01, ***p < 0.001, ****p < 0.0001, ns = not significant.

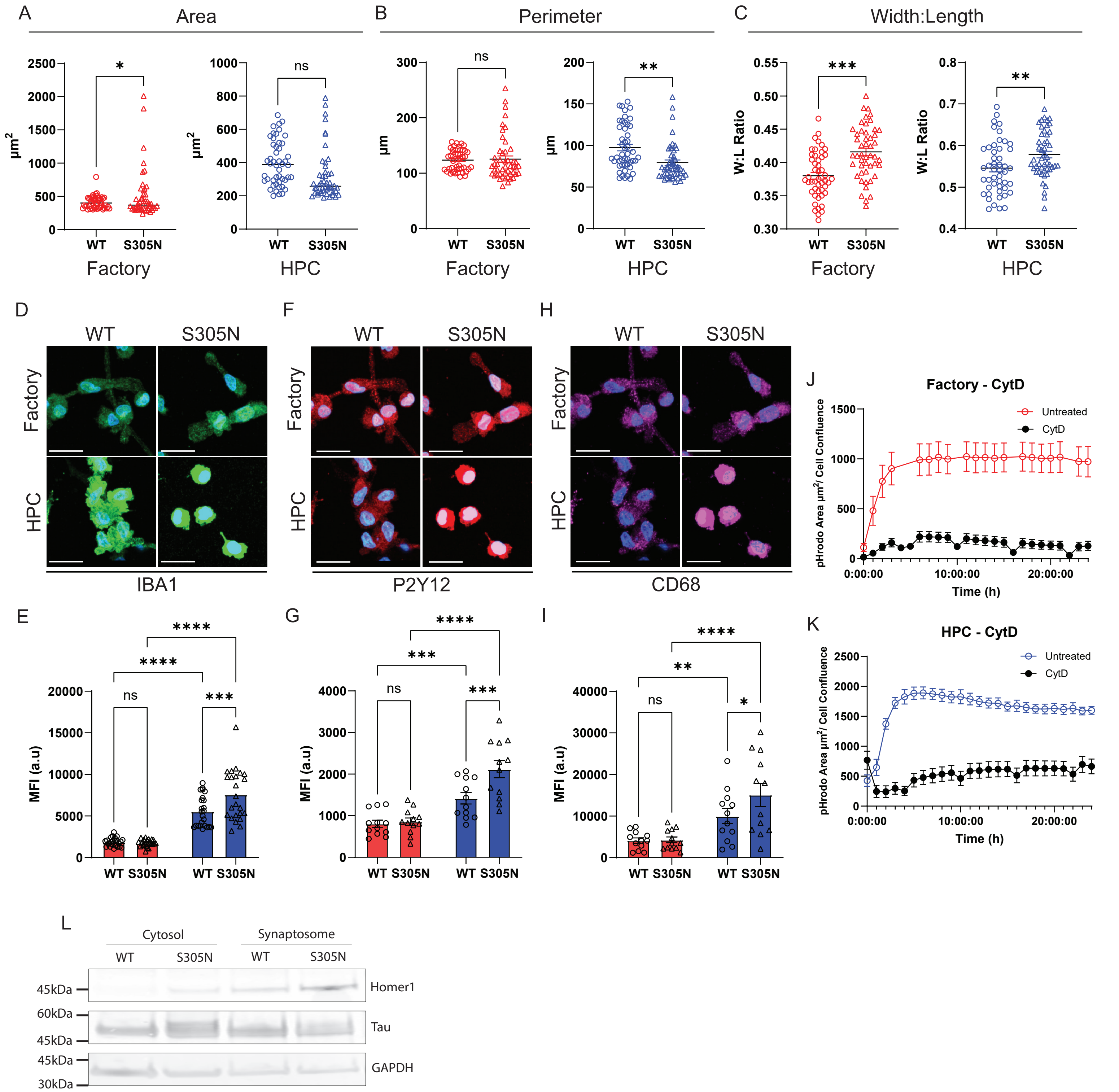


**Figure S2.** ***MAPT*-S305N causes mild morphological changes in microglia derived from both protocols. Related to Figure 2.**

**A - C.** Quantification of morpholigical features of Factory and HPC microglia dependent on MAPT genotype. n = 4 fields of view (FOV) from 4 independent wells per replicate. Data points represent mean value per FOV. Two-way ANOVA, Uncorrected Fisher's LSD.

**D-I** Representative images and quantification of mean fluorescence intensity (MFI) of ***D-E***. IBA1, ***F-G.*** P2Y12 and ***H-I.*** CD68 in WT and *MAPT*-S305N microglia derived from either Factrory or HPC protocols. Scale bar = 20µm.. n = 4 fields of view (FOV) from 2 independent wells per replicate. Data points represent mean value per FOV. Mixed effects analysis, Tukey's multiple comparisons test.

**J-K**. Quantification of pHrodo positive area normalised to cell confluence over 24 hours following E.Coli exposure with and without cytochalasin D (CytD) in ***J.*** Factory and ***K.*** HPC microglia.

**L.** Validation of synaptosome isolation from cortical organoids.

All data derived from N = 2 independent isogenic pairs of iPSC lines, 3 independent differentiations per protocol. Error bars = SEM. *p < 0.05, **p < 0.01, ***p < 0.001, ****p < 0.0001, ns = not significant.

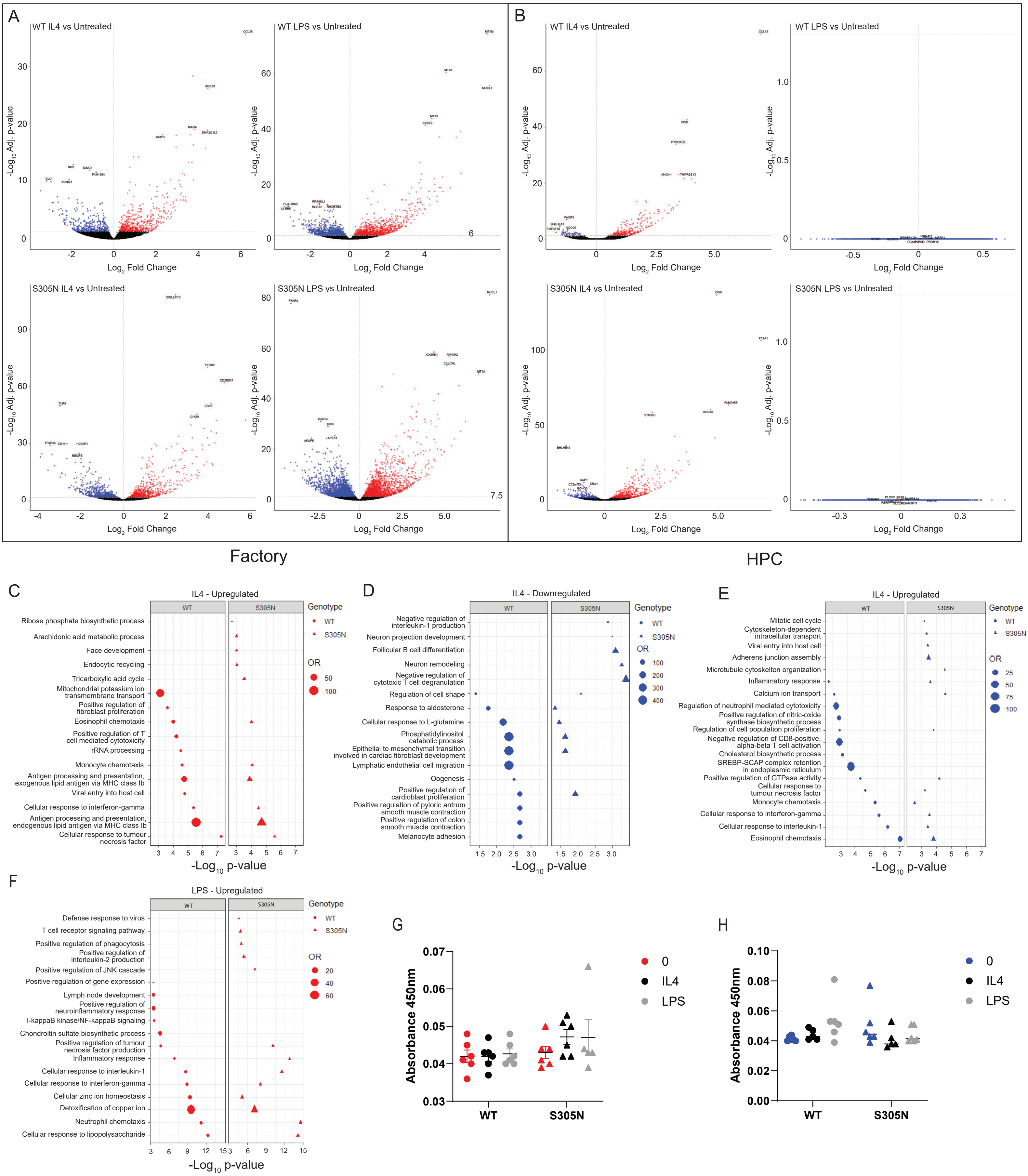


**Figure S3. Factory and HPC protocol microglia have distinct responses to inflammatory stimuli. Related to Figure 3.**

**A-B.** Differentially expressed genes in ***A.*** Factory and ***B.*** HPC microglia following LPS and IL4 stimulation, separated by *MAPT* genotype.

**C.** Upregulated pathways in Factory microglia following IL4 stimulation.

**D-E.** Pathway emrichment of ***D.*** downregulated and ***E.*** upregulated differentially expressed genes n HPC microglia following IL4 stimulation.

***F.*** Upregulated pathways in Factory microglia following LPS stimulation.

**G-H.** Absorbance valies (450nm) for IL21 ELISA on conditioned media from ***G.*** Factory and ***H.*** HPC-derived microglia

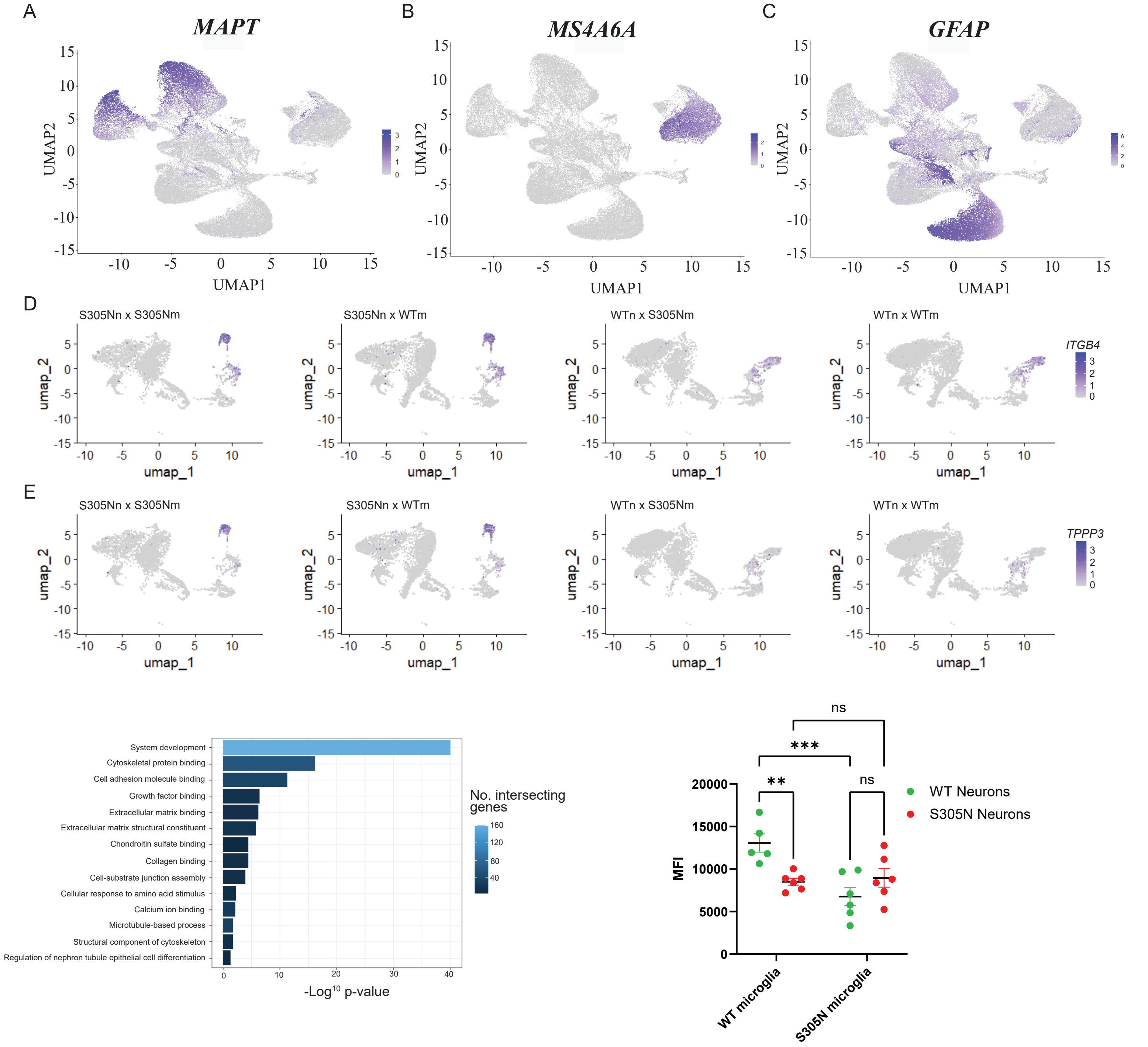


**Figure S4. A population of microglia from both genotypes are capable of resonding to *MAPT* mutation neurons, but *MAPT*-S305N microglia show impaired TMEM176B expression. Related to Figure 4.**

**A-C.** UMAP projection of tri-cultures following single-cell sequencing, with clusters identified as ***A.*** Neurons (*MAPT* expression), ***B.*** Microglia (*MS4A6A* expression) and C. Astrocytes (*GFAP* expression). **D-E.** UMAP projection of iPSC microglia under different co-culture conditions, highlighting ***D.*** *ITGB4* and ***E.*** *TPPP3* expression in clusters unique to *MAPT*-S305N neuron co-culture. **F.** Pathway enrichment of significantly upregulated marker genes in unique cluster highlighted in ***D-E***. Sequencing data derived from N = 2 independent isogenic pairs of iPSC lines from one round of differentiation (n = 24,197 microglia, 34,962 neurons). F. Mean fluorescence intensity (MFI) of TMEM176B in IBA1 positive cells in different genotype co-culture conditions. N = 1 isogenic pair of iPSC lines from three independent rounds of differentiation, with 2 replicate wells analysed per batch. Two-way ANOVA, uncorrected Fisher's LSD multiple comparisons test. Error bars = SEM. $^{**}p < 0.01$, $^{***}p < 0.001$, ns = not significant.
